# Intrathecal catheter implantation decreases cerebrospinal fluid dynamics in cynomolgus monkeys

**DOI:** 10.1101/2020.07.27.222646

**Authors:** Mohammadreza Khani, Audrey Q. Fu, Joshua Pluid, Christina P. Gibbs, John N. Oshinski, Tao Xing, Gregory R. Stewart, Jillynne R. Zeller, Bryn A. Martin

**Affiliations:** Department of Chemical and Biological Engineering, University of Idaho, Moscow, ID, USA; Department of Statistical Science, University of Idaho, Moscow, ID, USA; Department of Radiology and Imaging Sciences, Emory University, Atlanta, GA, USA; Department of Mechanical Engineering, University of Idaho, Moscow, ID, USA; Axovant, New York, NY, USA; Voyager Therapeutics, Cambridge, MA, USA; Northern Biomedical Research, Spring Lake, MI, USA

## Abstract

A detailed understanding of the CSF dynamics is essential for testing and evaluation of intrathecal drug delivery. Preclinical work using large-animal models (e.g., monkeys, dogs and sheep) has great utility for defining spinal drug distribution/pharmacokinetics and provide an important tool for defining safety. In this study, we investigated the impact of catheter implantation in the sub-dural space on CSF flow dynamics in Cynomolgus monkeys. Magnetic resonance imaging (MRI) was performed before and after catheter implantation to quantify the differences based on catheter placement location in the cervical compared to the lumbar spine. Several geometric and hydrodynamic parameters were calculated based on the 3D segmentation and flow analysis. Hagen-Poiseuille equation was used to investigate the impact of catheter implantation on flow reduction and hydraulic resistance. A linear mixed-effects model was used in this study to investigate if there is a statistically significant difference between cervical and lumbar implantation, or between two MRI time points. Results showed that geometric parameters did not change statistically across MRI measurement time points and did not depend on catheter location. However, catheter insertion did have a significant impact on the hydrodynamic parameters and the effect was greater with the cervical implantation. CSF flow rate decreased up to 54.7% when the catheter located in the cervical region. The maximum flow rate reduction in the lumbar implantation group was 21%. Overall, lumbar catheter implantation disrupted CSF dynamics to a lesser degree than cervical catheter implantation and this effect remained up to two weeks post-catheter implantation

## Background and Introduction

A detailed understanding of the CSF dynamics is needed for testing and evaluation of intrathecal drug delivery associated with catheter insertion. The cerebrospinal fluid (CSF) is secreted from arterial blood by the choroid plexus of the lateral and fourth ventricles by a combined process of diffusion, pinocytosis and active transfer (1–3). A small amount is produced by ependymal cells. The circulation of CSF is aided by the pulsations of the choroid plexus and by the motion of the cilia of ependymal cells (4, 5). CSF is absorbed across the arachnoid villi into the venous circulation and a significant amount probably also drains into lymphatic vessels around the cranial cavity and spinal canal (6, 7). CSF acts as a cushion that protects the brain from mechanical insult and supports the venous sinuses (5). It also plays an important role in the homeostasis and metabolism of the central nervous system (8).

In intrathecal drug delivery, medications are introduced directly to the spinal fluid (intrathecal space) through a drug delivery system. An externalized intrathecal catheter is the most widely used technique for administration of intrathecal drugs (9). With intrathecal delivery, less medication is necessary than if the medication was taken orally, and fewer side effects are often seen. Currently approved medications for intrathecal administration by the U.S. Food and Drug Administration (FDA) include morphine, ziconotide and baclofen. For these therapies, the doctor places a catheter beneath the skin and into the space along the spine (the intrathecal space) to release the drug into the cerebrospinal fluid. With intrathecal delivery, the drug can bypass the blood-brain barrier and more directly reach nervous system tissue. If the goal of treatment is to reduce spasticity in both the arms and the legs, an intrathecal catheter can be placed in a more rostral position potentially leading to increased uniformity in baclofen dosing of the cervical and lumbar spine and improved reduction in spasticity of the upper and lower limbs (10).

Preclinical studies need to be performed using large-animal models (e.g., monkeys, dogs and sheep) since these models have utility to define spinal drug distribution/pharmacokinetics and provide an important tool for assessment of drug safety. Preclinical studies also provide insight into the potential mechanisms of intrathecal drug delivery. Cynomolgus monkeys are a commonly used animal model for these studies because of their similarity to humans with regard to the pathophysiology of a variety of diseases and presumed similarity with regard to central nervous system (CNS) anatomy and CSF hydrodynamics.

There have been previous reports of CSF analyses in nonhuman primates. Acute procedures include cisterna magna tap in anesthetized rhesus monkeys and baboons (11) and lumbar puncture in anesthetized rhesus monkeys (12), baboons(13), and chimpanzees (14). Chronic procedures include lumbar puncture with needle and stylet, where the needle remains in place for periods of 90 to 300 minutes (15). Taylor et al. reported cannulating the lumbar subarachnoid space (SAS), which allowed CSF collection from a rhesus monkey for 72 hours (16). Perlow catheterized the SAS of a rhesus monkey by inserting a catheter into the lumbar region and advancing it cephalad so that the tip terminated in the cisternal-cervical SAS (17). CSF was then withdrawn continuously by a peristaltic pump for 48 hours. However, these reports provided relatively few details of the spinal tap procedures nor specifications of the apparatus, such as the gauge of the cannula or catheter. Also, none of these procedures continued for more than 48 hours. Thus, the potential impact of prolonged intrathecal catheterization on CSF dynamics was not analyzed.

To our knowledge, no studies have investigated how catheter placement may impact CSF dynamics in Cynomolgus monkeys. Our previous study developed a quantitative method to characterize CSF dynamics and geometry in non-human primates (NHPs) (18). This method was demonstrated to reliably measure CSF dynamics parameters over a two-week period in a group of eight NHPs. The goal of the current study was to apply the same MRI measurements and post-processing methods on a series of scans collected for the same cohort of NHPs to quantify: a) alterations in CSF dynamics due to catheter placement in the intrathecal space, b) track these changes over time, and c) determine if there are any differences that occur based on catheter placement location in the cervical compared to lumbar spine.

## Materials and methods

### Ethics statement

This study was submitted to and approved by the local governing Institutional Animal Care and Use Committee at Northern Biomedical Research (IACUC approval #084-014A, Spring Lake, MI). This study did not unnecessarily duplicate previous experiments and alternatives to the use of live animals were considered. Procedures used in this study were designed with consideration of the well-being of the animals.

### Catheter Placement and Parameters

Eight (NHP 01-08) healthy cynomolgus monkeys (Macaca fascicularis, origin Mauritius) were obtained from Charles River Research Models, Houston TX with an average weight of 4.4 ± 1.2 kg and age of 4.6 ± 0.4 years (mean ± standard deviation) (**Table 1**). NHP 01 was male and all other NHPs were female (02-08). These animals were purpose-bred and experimentally naïve.

**Table 1.** Cynomolgus monkey case information.

| Designation | Gender | Catheter Placement | Weight (kg) | Age (yr.) |
| --- | --- | --- | --- | --- |
| NHP 01 | M | C5 | 4.0 | 4.1 |
| NHP 02 | F | C5 | 3.3 | 4.7 |
| NHP 03 | F | L1 | 5.1 | 4.5 |
| NHP 04 | F | L1 | 3.2 | 4.4 |
| NHP 05 | F | C5 | 4.8 | 4.2 |
| NHP 06 | F | C5 | 3.0 | 4.9 |
| NHP 07 | F | L1 | 4.2 | 5.2 |
| NHP 08 | F | L1 | 5.8 | 4.4 |

Each NHP was scanned with an identical MRI protocol (see MRI methods) across all study time points (**Fig 1**). MRI_PRE-1_ and MRI_PRE-2_ were spaced 14 days apart prior to catheter placement. MRI_PRE-1_ and MRI_PRE-2_ were used in our previous publication (19) to quantify reliability of CSF flow parameters. At day 17, the NHP’s were randomly assigned to have intrathecal catheter implantation (IT-PEPU-35, SAI Infusion Technologies, Lake Villa, IL, U.S.A.) in the spinal SAS at C5 (Cervical Group, n = 4) or L1 (Lumbar Group, n = 4). The catheter had the following dimensions: first 10 cm distal to the tip (ID = 0.38 mm and OD = 0.99 mm), next 24 cm (ID = 1.19 mm and OD = 1.98 mm) and last ~1.5 cm (ID = 1.07 mm and OD = 1.93 mm). Implantation was performed by fluoroscopic imaging with contrast agent. Catheter patency was verified by visual inspection and confirmation of the ability to withdraw CSF from the port/catheter system at day 24 and 28. MRI_POST-1_ was collected on day 31 to determine the acute impact of implantation on CSF dynamics and geometry by comparison of results to MRI_PRE-2_. MRI_POST-2_ was collected at day 45 to determine if this impact persisted after implantation.

**Fig 1.**
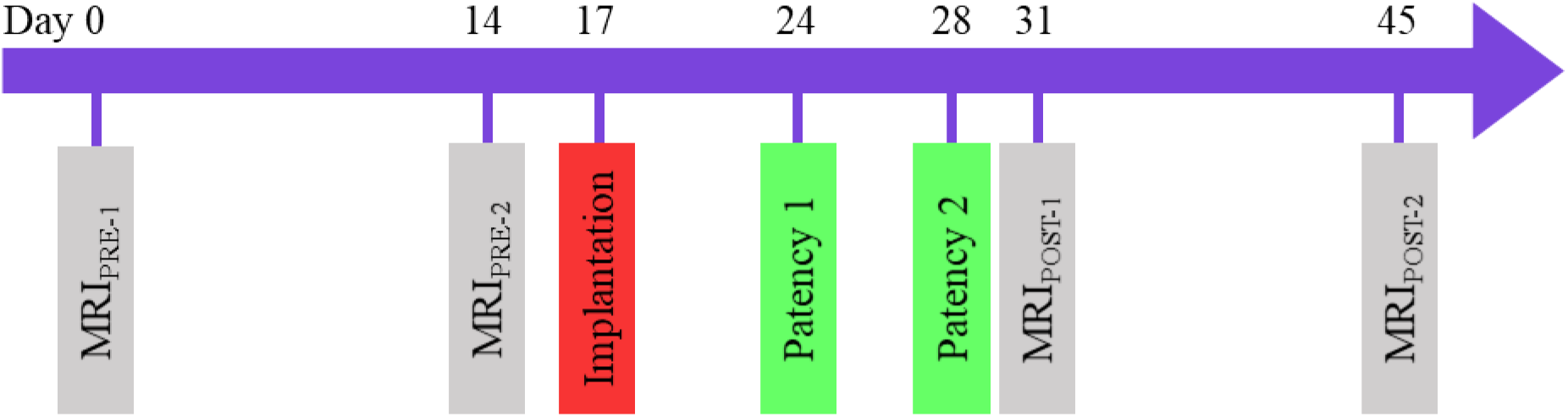
Study outline. This study involved a total of four MRI scans collected over a period of 45 days. MRI_PRE-1_ and MRI_PRE-2_ were performed prior to catheter implantation to confirm scan consistency (see results published in Khani et al. (19)). An intrathecal catheter was implanted within the cervical SAS (C5, n=4) and lumbar SAS (L1, n=4). Catheter patency was confirmed on day 24 and 28. MRI_POST-1_ was collected to determine the acute impact of catheter implantation compared to MRI_PRE-2_. MRI_POST-2_ was collected to determine if this impact persisted two weeks after implantation.

### MRI scan protocols

MRI scan protocols were previously described in detail by Khani et al. (18). In brief, all MRI measurements were acquired at Northern Biomedical Research (Norton Shores, Michigan, U.S.A.) on a Philips 3T scanner (Achieva, software V2.6.3.7, Best, The Netherlands). Prior to MRI scanning each NHP was prepared using standard procedures and precautions. NHPs were positioned in the scanner in the supine position without assistance from artificial respiration. During each scan, heart rate and respiration were monitored continuously with ~ 1 liter/minute of oxygen and 1-3% isoflurane anesthetic administered via an endotracheal tube for sedation.

A stack of high-resolution axial T2-weighted MR images of the complete spinal SAS geometry was acquired for each NHP. The anatomical region scanned was ~30 cm in length, which included the intrathecal SAS below the lower brain stem extending caudally to the filum terminale. Thru-plane (head-foot, z-direction) CSF flow was measured by phase-contrast MRI (PC-MRI) images collected at six axial locations along the spine for each NHP. Axial locations were marked at the foramen magnum (FM), C2-C3, C5-C6, T4-T5, T10-T11, and L3-L4. The slice location for each scan was oriented approximately perpendicular to the CSF flow direction with slice planes intersecting vertebral discs (**Fig 2**).

**Fig 2.**
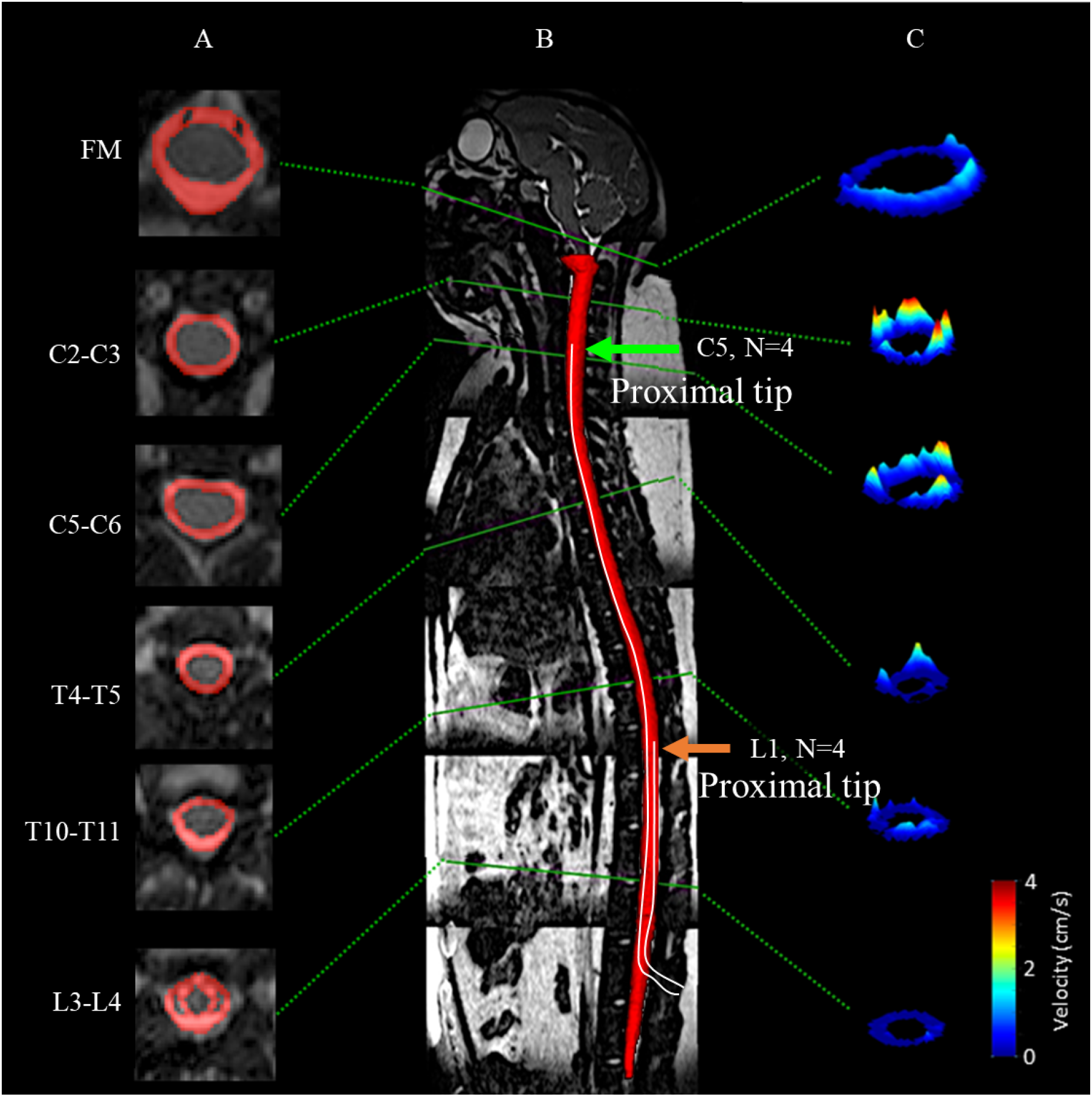
Manual segmentation of the spinal SAS using a T2-weighted MR image and axial PC-MRI and CSF velocity profiles at corresponding vertebral levels for a cynomolgus monkey analyzed in this study. (A) Visualization of SAS area manually selected around the spinal cord at multiple axial levels. (B) Mid-sagittal high-resolution T2-weighted MRI and 3D visualization of entire SAS geometry. (C) 3D visualization of peak systolic CSF velocity profiles based on in vivo PC-MRI measurements at FM, C2-C3, C5-C6, T4-T5, T11-T12, and L3-L4. Arrows represent the location of catheter placement at the cervical (C5) or lumbar (L1) implantation groups (N=4 NHPs in each group).

### Image segmentation and flow analysis

The high-resolution T2-weighted anatomic MRI images were semi-automatically segmented using the free open-source ITK-snap software (Version 3.0.0, University of Pennsylvania, U.S.A.) (20), which provided semi-automatic segmentation using active contour methods, as well as manual delineation and image navigation (**Fig 2A**). The manual segmentation tool was used most frequently with the view of the three orthogonal planes. The catheter was considered to be an empty region within the spinal SAS, because it was not possible to consistently identify within the MR images due to its small lumen diameter. Once the segmentation was complete, the 3D model (**Fig 2B**) was exported in a .STL (Stereo Lithography) format for subsequent analysis as outlined below. Detailed information on the segmentation procedure is provided by Khani et al. (18).

CSF flow was quantified at six axial locations along the spine (**Fig 2C**) using GTFLOW software (64-bit, Version 2.2.10, Gyrotools, Zurich, Switzerland) by the procedure previously described by Khani et al. (18). The six distinct flow rates were smoothed in a spatial-temporal fashion using MATLAB and a 2D “fit” function with the fit-type designated as “smoothing-spline”. Since heart rate variability was present between the PC-MRI scans, the CSF flow waveform timing was normalized to the average heart rate for all NHPs. An average spatial-temporal CSF waveform was determined for each case. CSF pulse wave velocity, *PWV*, was computed based on the slope of the arrival time of peak CSF flow along the spine (21).

### Geometric and hydrodynamic parameter quantification

Several geometric and hydrodynamic parameters were calculated based on the 3D segmentation and flow analysis using our previously published methods (18). Total SAS surface area, *SA*_*sas*_, was calculated as the sum of the surface area of spinal cord, *SA*_*c*_, and dura, *SA*_*d*_. Spinal cord nerve roots were not included in the surface area calculation of the cord since these small features were not possible to accurately visualize by MR imaging. Total volume of the SAS, *V*_*sas*_, was computed by subtracting the volume of the spinal cord, *V*_*c*_ from the volume of the dura, *V*_*d*_. Total SAS length, *L*_*sas*_, from the FM to the SAS termination was quantified.

Axial distribution of the SAS cross-sectional area, *A*_*sas*_ (*z*), was based on cross-sectional area of the spinal cord at that location, *A*_*c*_ (*z*), and dura, *A*_*d*_ (*z*). The axial distribution of the catheter cross-sectional area for the lumbar and cervical catheters were subtracted for MRI_POST-1_ and MRI_POST-2_. Similarly, hydraulic diameter, *D*_*h*_ (*z*) = 4 *A*_*sas*_ (*z*) / *P*_*sas*_ (*z*), was determined based on the wetted perimeter, *P*_*sas*_ (*z*), with the perimeter computed as the sum of the spinal cord, *P*_*c*_ (*z*), and dura, *P*_*d*_ (*z*), perimeters at each z-location. The axial distribution of the catheter perimeter for the lumbar and cervical catheters were added for MRI_POST-1_ and MRI_POST-2_. Axial distribution of CSF stroke volume was computed as *SV*(*z*) = ∫|*Q*(*z*,*t*)|*dt*, where |*Q*(*z*, *t*)| is the absolute value (22). Peak systolic (toward feet) and diastolic (toward the head) CSF flow rate was quantified as *Q*_*sys*_ (*z*) and *Q*_*dia*_ (*z*), and the CSF flow rate amplitude was given by *Q*_*a*_ (*z*) = *Q*_*dia*_ (*z*) = *Q*_*sys*_ (*z*). Spatial mean thru-plane velocity at peak systole was computed as 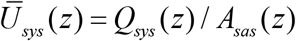 and at diastole as 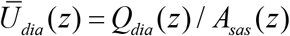. Reynolds number was computed as 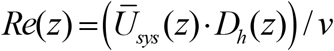, where *v* is the kinematic viscosity of CSF at body temperature, 0.693 mPa·s (23). Womersley number was computed as 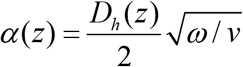, where *ω* is the angular velocity (*ω*= 2*π*/ *T*) of the volume flow waveform with *T* equal to the heart rate. To allow parameter comparison across NHPs, each parameter’s axial distribution for each NHP was normalized to the average *L*_*sas*_ measured for all NHPs. After normalization, the mean axial distribution for each parameter was computed for each group (Cervical or Lumbar catheter implantation) at each MRI time point (MRI_PRE-2_, MRI_POST-1_ and MRI_POST-2_).

Catheter implantation could potentially reduce CSF flow due to increased hydraulic resistance. We estimated the flow reduction for the cervical and lumbar implantation group by: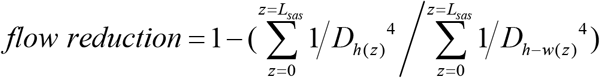. Where *D*_*h*_*(z)*, is the axial distribution of hydraulic diameter for MRI_PRE-2_, and *D*_*h-w*_*(z)*, is the predicted hydraulic diameter by taking into account the axial distribution of catheter area and perimeter for the lumbar and cervical groups. This flow reduction is approximated based on the Hagen-Poiseuille equation for steady, incompressible, laminar pipe flow under the assumption that intracranial pressure pulsations, that drive CSF flow along the spine, are not affected by presence or absence of the catheter (i.e. *Δp* = constant).

### Statistical analysis

We hypothesized that implantation of the catheter would decrease CSF dynamics and geometry, and that these changes would be elevated for NHPs with cervical implantation compared to lumbar implantation. For each of the parameters investigated here, it was measured at multiple locations along the spinal cord for each cynomolgus monkey and MRI measurement with a certain implantation. Since the NHPs were randomly selected from a population, we developed the following linear mixed-effects model:

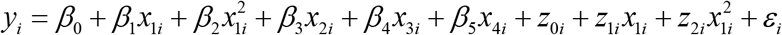

Where *y*_*i*_ is the parameter of interest (a geometric or hydrodynamic parameter), *x*_1*i*_ is the location, *x*_2*i*_ is the catheter location (cervical / lumbar) or the MRI time point (e.g., MRI_PRE-2_, MRI_POST-1_ and MRI_POST-2_), *x*_3*i*_ and *x*_4*i*_ are the age and weight of each NHP, respectively. The random error, *ε*_*i*_, has a normal distribution with mean 0 and variance σ2: *ε*_*i*_~*N*(0,σ^2^). While *β* are fixed effect sizes, *z* represent the random-effect coefficients, which follow a multivariate normal distribution with mean of 0 and a symmetric variance-covariance matrix:

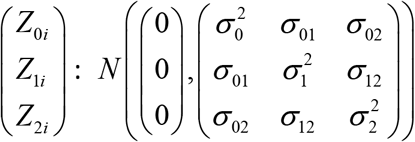

We used the “*fitlme*” function in Matlab (Ver. R2019a Mathworks Corp., Natick, MA) to estimate the parameters in this linear mixed-effects model and test the hypothesis.

This model treats the catheter location and the MRI time point as a fixed effect, with the corresponding coefficient indicating the effect size. We could further test whether the true effect size is significantly different from zero. If so, it means that there is a statistically significant difference between cervical and lumbar implantation, or between two MRI time points. This model treats the NHPs as random; this means that the multiple measurements from an NHP can form a curve, and that this curve may be different from one NHP to another.

Using this linear mixed-effects model, we estimated the relative effect sizes of the following seven pairs: four pairs comparing time points (time points PRE-2C versus POST-1C for cervical implantation; time points PRE-2L versus POST-1L for lumbar implantation; time points PRE-2C versus POST-2C for cervical implantation; times points PRE-2L versus POST-2L for lumbar implantation), and three comparing cervical versus lumbar implantation (at time point PRE-2, POST-1 and POST-2). For each pair, we tested the statistical significance of the two groups being different and obtained a P value. Since we performed this analysis for 13 geometric and hydrodynamic parameters, we derived 13 × 7 = 91 P values. Many of these P values were dependent due to the strong dependence among several parameters of interest. We accounted for multiple comparison with Bonferroni correction by adjusting the threshold for P values to be 0.05/91=5.49e-4. This identified a highly conservative set of significant P values. Note that this approach assumes independence among P values. When two parameters of interest are highly correlated, they would lead to similar P values that are both identified to be significant after correction. In this case, we can only conclude that one or both parameters are significant, but we cannot pinpoint the truly significant parameter.

## Results

A summary of geometric and hydrodynamic parameter results obtained at MRI_PRE-2_, MRI_POST-1_, and MRI_POST-2_ for the cervical (C) and lumbar (L) groups are shown in **Table 2** (Mean ± STD). Statistical assessments revealed that multiple hydrodynamic parameters were statistically different across study groups and time points (**Table 3**). However, geometric parameters were largely unchanged.

**Table 2.** Cynomolgus monkey geometric and hydrodynamic parameter results at each measurement time point and for the cervical and lumbar implantation groups. Note: The mean axial distribution for each parameter is shown based on N=4 NHPs in each group.

| Parameters | | MRI<sub>PRE-2C</sub><br>Mean ± STD | | MRI<sub>PRE-2L</sub><br>Mean ± STD | | MRI<sub>POST-1C</sub><br>Mean ± STD | | MRI<sub>POST-1L</sub><br>Mean ± STD | | MRI<sub>POST-2C</sub><br>Mean ± STD | | MRI<sub>POST-2L</sub><br>Mean ± STD | |
| --- | --- | --- | --- | --- | --- | --- | --- | --- | --- | --- | --- | --- | --- |
| Geometric parameter<br>(Mean value along spine) | $P_c$ (mm) | 13.39 | 2.06 | 14.06 | 1.64 | 13.31 | 2.10 | 14.09 | 1.44 | 13.79 | 2.13 | 14.65 | 1.20 |
| | $P_d$ (mm) | 21.53 | 1.96 | 22.30 | 1.66 | 21.81 | 1.91 | 22.30 | 1.47 | 22.05 | 1.97 | 22.55 | 1.38 |
| | $P_{sas}$ (mm) | 35.72 | 4.38 | 37.46 | 3.91 | 39.29 | 4.15 | 38.06 | 3.08 | 40.60 | 4.82 | 39.02 | 2.88 |
| | $A_c$ (mm<sup>2</sup>) | 14.40 | 2.72 | 15.47 | 3.07 | 14.89 | 3.25 | 16.12 | 2.57 | 15.46 | 3.46 | 16.24 | 2.25 |
| | $A_d$ (mm<sup>2</sup>) | 38.21 | 6.22 | 40.10 | 5.89 | 38.89 | 6.00 | 40.05 | 5.11 | 39.91 | 6.05 | 40.72 | 4.88 |
| | $A_{sas}$ (mm<sup>2</sup>) | 23.81 | 4.66 | 24.63 | 4.31 | 22.51 | 4.50 | 23.76 | 3.76 | 22.96 | 4.17 | 24.30 | 3.62 |
| | $SA_c$ (cm<sup>2</sup>) | 40.24 | 3.38 | 42.40 | 2.71 | 38.68 | 1.67 | 41.32 | 2.80 | 39.85 | 3.43 | 42.97 | 1.76 |
| | $SA_d$ (cm<sup>2</sup>) | 64.85 | 3.83 | 67.37 | 3.54 | 63.59 | 4.72 | 65.55 | 3.03 | 63.87 | 4.90 | 66.32 | 2.47 |
| | $SA_{sas}$ (cm<sup>2</sup>) | 105.09 | 7.17 | 109.78 | 6.07 | 102.27 | 6.12 | 106.87 | 5.74 | 103.72 | 8.22 | 109.29 | 4.22 |
| | $V_c$ (mL) | 4.33 | 0.39 | 4.70 | 0.59 | 4.35 | 0.41 | 4.76 | 0.44 | 4.48 | 0.39 | 4.80 | 0.32 |
| | $V_d$ (mL) | 11.51 | 1.18 | 12.13 | 1.21 | 11.35 | 1.10 | 11.80 | 0.95 | 11.59 | 1.03 | 12.00 | 0.87 |
| | $V_{sas}$ (mL) | 7.17 | 0.82 | 7.43 | 0.79 | 6.57 | 0.77 | 6.99 | 0.57 | 6.68 | 0.67 | 7.15 | 0.58 |
| Hydrodynamic parameter<br>(Mean value along spine) | $D_h$ (mm) | 2.67 | 0.34 | 2.66 | 0.37 | 2.35 | 0.43 | 2.54 | 0.34 | 2.31 | 0.29 | 2.52 | 0.29 |
| | $Re$ | 32.32 | 14.12 | 28.51 | 13.12 | 15.13 | 11.74 | 21.77 | 11.61 | 10.79 | 6.58 | 21.45 | 13.03 |
| | $\alpha$ | 5.45 | 0.86 | 5.43 | 0.65 | 4.61 | 1.04 | 5.05 | 0.66 | 4.42 | 0.56 | 5.09 | 0.59 |
| | $U_{peak-sys}$ (cm/s) | -0.92 | 0.42 | -0.84 | 0.36 | -0.49 | 0.40 | -0.63 | 0.32 | -0.34 | 0.21 | -0.64 | 0.39 |
| | $U_{peak-dia}$ (cm/s) | 0.66 | 0.30 | 0.56 | 0.23 | 0.40 | 0.29 | 0.46 | 0.23 | 0.32 | 0.19 | 0.49 | 0.26 |
| | $Q_{peak-sys}$ (mL/s) | -0.21 | 0.09 | -0.19 | 0.08 | -0.11 | 0.08 | -0.15 | 0.08 | -0.08 | 0.04 | -0.15 | 0.09 |
| | $Q_{peak-dia}$ (mL/s) | 0.15 | 0.06 | 0.13 | 0.05 | 0.08 | 0.05 | 0.11 | 0.05 | 0.07 | 0.04 | 0.11 | 0.06 |
| | $Q_a$ (mL/s) | 0.36 | 0.22 | 0.32 | 0.25 | 0.19 | 0.11 | 0.25 | 0.20 | 0.15 | 0.08 | 0.26 | 0.21 |
| | $SV$ (cm<sup>3</sup>) | 0.06 | 0.02 | 0.05 | 0.03 | 0.03 | 0.02 | 0.05 | 0.03 | 0.03 | 0.01 | 0.04 | 0.03 |
| | $PWV$ (cm/s) | 1.15 | 1.21 | 1.11 | 0.21 | 1.25 | 0.59 | 1.16 | 0.52 | 1.16 | 0.06 | 1.09 | 0.21 |

**Table 3.** Statistical comparison of parameters across measurement time points for baseline vs. follow-up MRIs and cervical vs. lumbar catheter insertion. P values are obtained from linear mixed effects model (see “Statistical analysis” section for details).

| Parameters |  | Baseline vs Follow-up |  |  |  | Cervical vs Lumbar |  |  |
| --- | --- | --- | --- | --- | --- | --- | --- | --- |
|  |  | MRI<sub>PRE-2C</sub><br>VS<br>MRI<sub>POST-1C</sub> | MRI<sub>PRE-2L</sub><br>VS<br>MRI<sub>POST-1L</sub> | MRI<sub>PRE-2C</sub><br>VS<br>MRI<sub>POST-2C</sub> | MRI<sub>PRE-2L</sub><br>VS<br>MRI<sub>POST-2L</sub> | MRI<sub>PRE-2C</sub><br>VS<br>MRI<sub>PRE-2L</sub> | MRI<sub>POST-1C</sub><br>VS<br>MRI<sub>POST-1L</sub> | MRI<sub>POST-2C</sub><br>VS<br>MRI<sub>POST-2L</sub> |
| Geometric | $A_d$ | 0.2042 | 0.9044 | 0.0027 | 0.2230 | 0.4912 | 0.3702 | 0.0157 |
| | $A_c$ | 0.0545 | 0.0247 | ** | 0.0056 | 0.0332 | *** | 0.3373 |
| | $A_{sas}$ | * | 0.0036 | 0.0175 | 0.2675 | 0.6254 | 0.3090 | 0.0612 |
| | $P_d$ | 0.0377 | 0.9536 | * | 0.0512 | 0.0170 | 0.3940 | 0.2902 |
| | $P_c$ | 0.6221 | 0.9015 | 0.0063 | ** | 0.3013 | 0.0055 | 0.5454 |
| | $P_{sas}$ | **** | 0.1155 | **** | *** | 0.0034 | 0.0012 | *** |
| Hydrodynamic | $D_h$ | **** | **** | **** | **** | 0.3255 | 0.5997 | 0.0080 |
| | $\alpha$ | **** | **** | **** | **** | 0.0565 | **** | 0.1960 |
| | $Re$ | **** | **** | **** | **** | **** | ** | **** |
| | $U_{peak-sys}$ | **** | **** | **** | **** | **** | 0.0873 | **** |
| | $U_{peak-dia}$ | **** | **** | **** | **** | 0.0299 | 0.0008 | **** |
| | $Q_a$ | **** | **** | **** | **** | **** | * | **** |
| | $SV$ | **** | **** | **** | **** | **** | 0.0026 | **** |

**P:** Probability value based on linear mixed effects model. The significance codes below use Bonferroni correction.
$p < 0.05/91 = *$ , $p < 0.01/91 = **$ , $p < 0.005/91 = ***$ , $p < 0.001/91 = ****$

### Geometric parameter results

Results indicated that cervical catheter insertion altered spinal SAS geometry to a greater degree than lumbar catheter insertion (**Fig 3**). Overall, 33 out of 42 geometric parameters did not change statistically across MRI measurement time points or depending on catheter location (**Table 3**).

**Fig 3.**
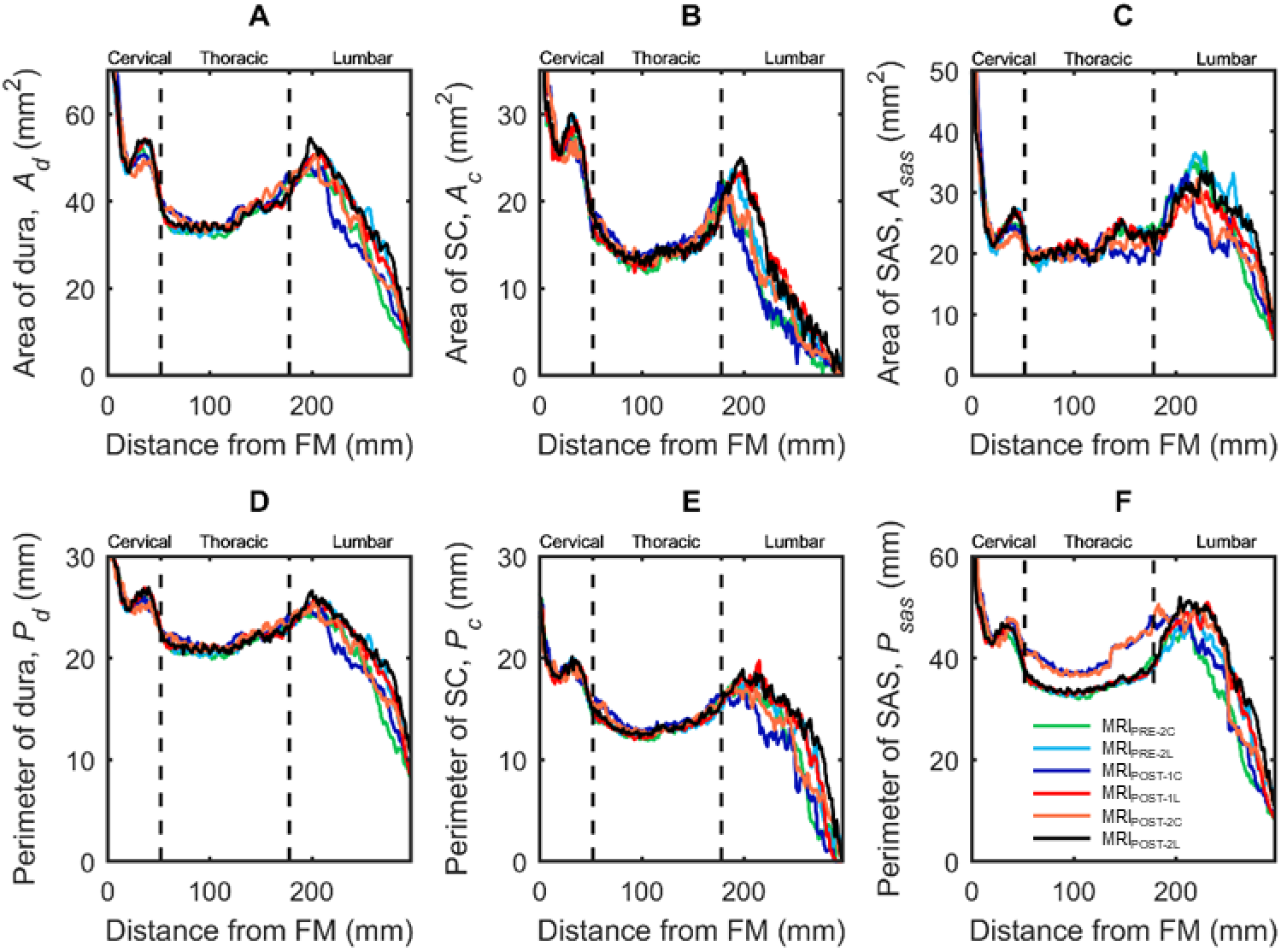
Axial distribution of geometric parameters computed along the spine for cynomolgus monkeys with cervical catheter implantation (MRI #C) or lumbar catheter implantation (MRI #L) measured prior to catheter implantation (MRI_PRE-2_), 17 days after catheter implantation (MRI_POST-1_), and 31 days after catheter implantation (MRI_POST-2_). (A) Area of dura, *A*_*d*_, (B) Area of spinal cord, *A*_*c*_, (C) Area of SAS, *A*_*sas*_, (D) Perimeter of dura, *P*_*d*_, (E) Perimeter of spinal cord, *P*_*c*_, (F) Perimeter of SAS, *P*_*sas*_. Each line corresponds to mean value of each NHPs group with catheter located in the lumbar (L) or cervical (c) spine before (MRI_PRE-2_) or after catheter placement (MRI_POST-1_ and MRI_POST-2_).

Axial distribution of geometric parameters showed relatively small changes across the lumbar and cervical implantation groups for *A*_*d*_, *A*_*c*_, *A*_*sas*_, *P*_*d*_, and *P*_*c*_ at all time points (**Fig 3A** through **E**). However, *P*_*sas*_ (**Fig 3F**) for the MRI_POST-1C_ and MRI_POST-2C_ groups increased significantly below the catheter tip after insertion (**Table 3**). Average CSF volume in the spinal SAS for all NHPs across all measurement time points (MRI_PRE-2_, MRIPOST_-1_ and MRI_POST-2_ for both cervical and lumbar groups) was 7.00 ml. Average cross-sectional area for spinal cord, dura and SAS for all NHPs was 15.43, 39.65 and 23.66 mm^2^, respectively. Average perimeter for spinal cord, dura and SAS was 13.88, 22.09, and 38.36 mm, respectively.

### Hydrodynamic parameter results

Catheter implantation was found to decrease CSF flow pulsations along the entire spine and this impact was greater for cervical catheter implantation compared to lumbar implantation (**Fig 4**). For example, MRI_POST-1C_ flow rate was lower than MRI_PRE-2C_ for all axial locations. Catheter implantation was found to decrease CSF flow pulsations even 31 days after catheter insertion (MRI_POST-2C_ and MRI_POST-2L_). These findings were supported by statistical analysis that showed changes in hydrodynamic parameters with cervical and lumbar catheter implantation to be highly significant for 40 out of 49 hydrodynamic parameters with p values < 0.05/91 (**Table 3**).

**Fig 4.**
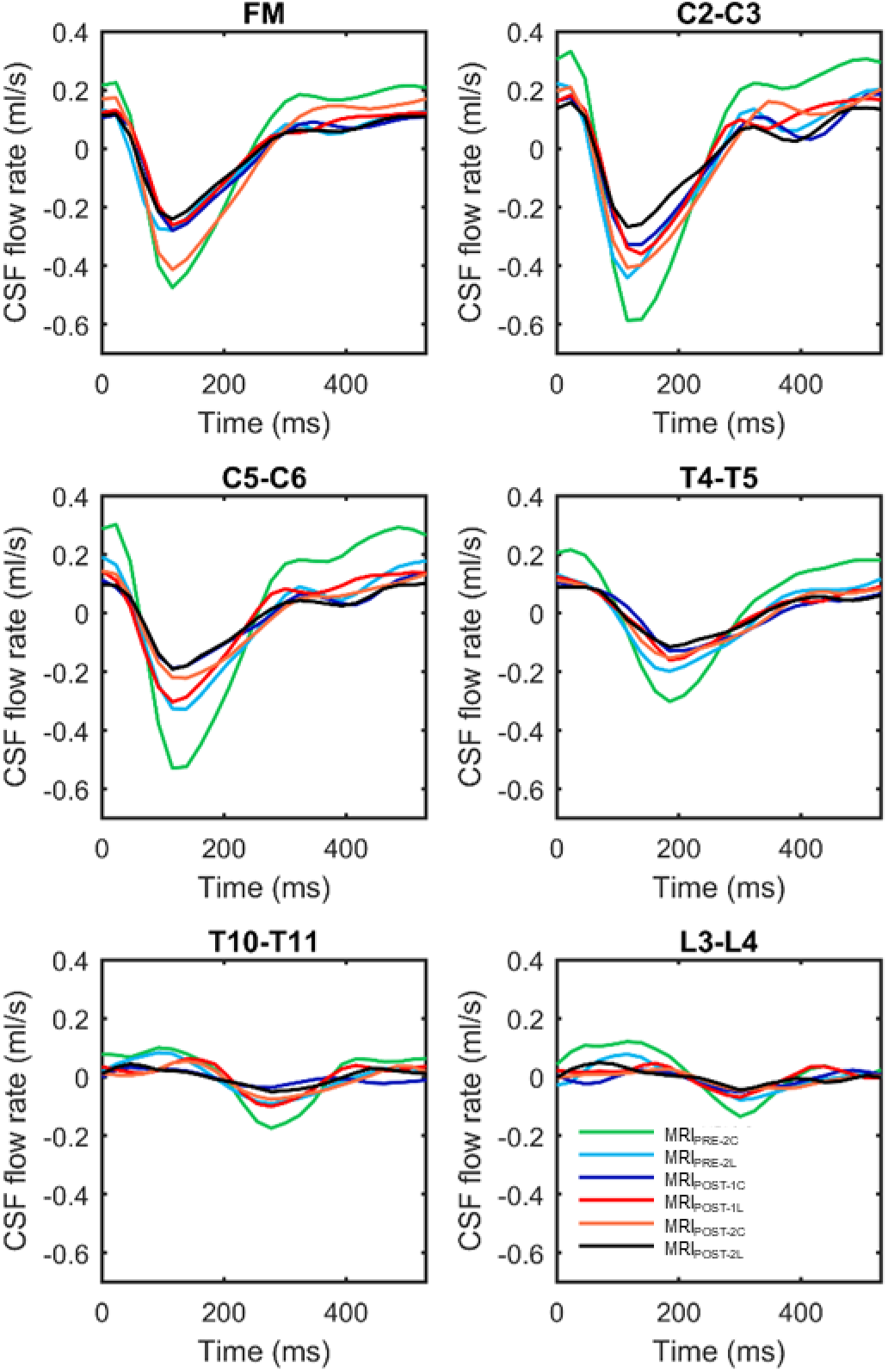
Average CSF flow waveforms for each MRI time point (4 NHPs at each point) measured at six axial locations along the spine (FM, C2-C3, C5-C6, T4-T5, T10-T11, L3-L4). Note: Peak systolic, CSF flow is in the caudal direction (negative values).

CSF flow rate of each NHP group quantified along the spine had a similar waveform shape, and axial distribution (**Fig 4**). CSF flow waveform showed a systolic peak at 100 to 150 ms in the cervical spine ranging from 0.2 - 0.6 (ml/s) for all NHPs. CSF flow rate at the C5-C6 for MRI_POST-1C_ and MRI_POST-2C_ was markedly smaller than both MRI_PRE-2C_ and _2L_, and MRI_POST-1L_ and MRI_POST-2L_ due to catheter placement within cervical SAS in those cases.

Average spatial-temporal distribution of the CSF flow along the spine showed a relatively smooth decrease in amplitude with a caudally directed CSF pulse wave velocity (**Fig 5**). Pulse wave velocity magnitude was similar across the groups and ranged from 1.09 – 1.24 m/s. Maximum CSF flow rate occurred for the MRI_PRE-2_ measurement within the cervical spine. Catheter placement decreased the flow rate spatially and temporally below the catheter tip in both MRI_POST-1_ and MRI_POST-2_.

**Fig 5.**
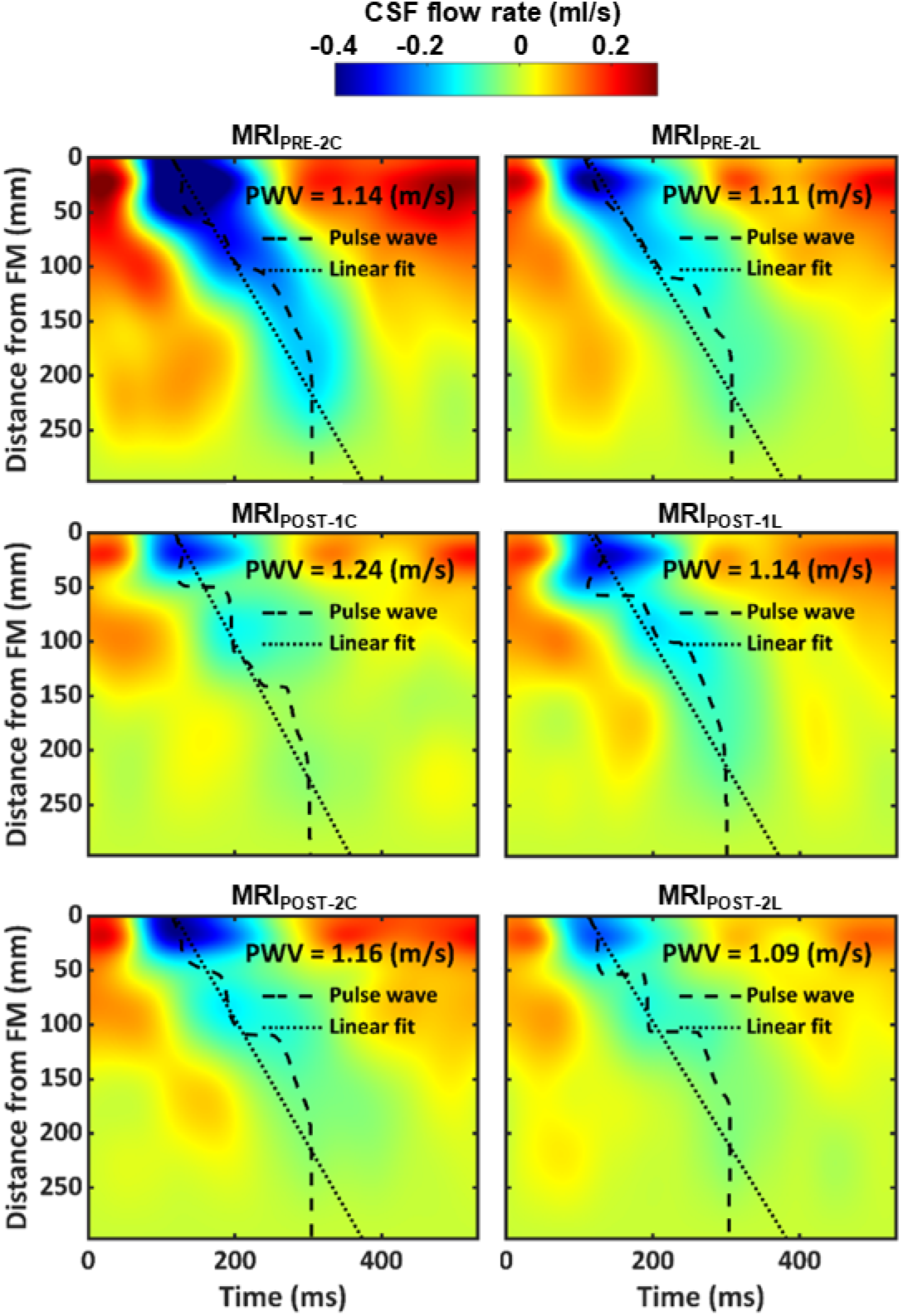
Mean CSF flow waveforms and Spatial-temporal distribution of CSF flow rate. Spatial-temporal distribution of the interpolated CSF flow rate along the spine for all cases measured by PC-MRI. Dashed line indicates peak CSF flow rate at each axial level and dotted line indicates linear fit on top of those values used to compute CSF pulse wave velocity (*PWV*).

Maximum *Re* number for MRI_PRE-2C_ was 80 at C3-C4 level (**Fig 6A**). MRI_POST-2C_ had the lowest *Re* value of 28 due to the cervical catheter implantation. Catheter implantation also decreased CSF flow rate amplitude (**Fig 6B**) and stroke volume (**Fig 6C**) at MRI_POST-1C_ and MRI_POST-2C_ compared to MRI_PRE-2C_ and for MRI_POST-1L_ and MRI_POST-2L_ compared to MRI_PRE-2L_. Albeit, the changes in flow rate amplitude and stroke volume were greater under cervical implantation.

**Fig 6.**
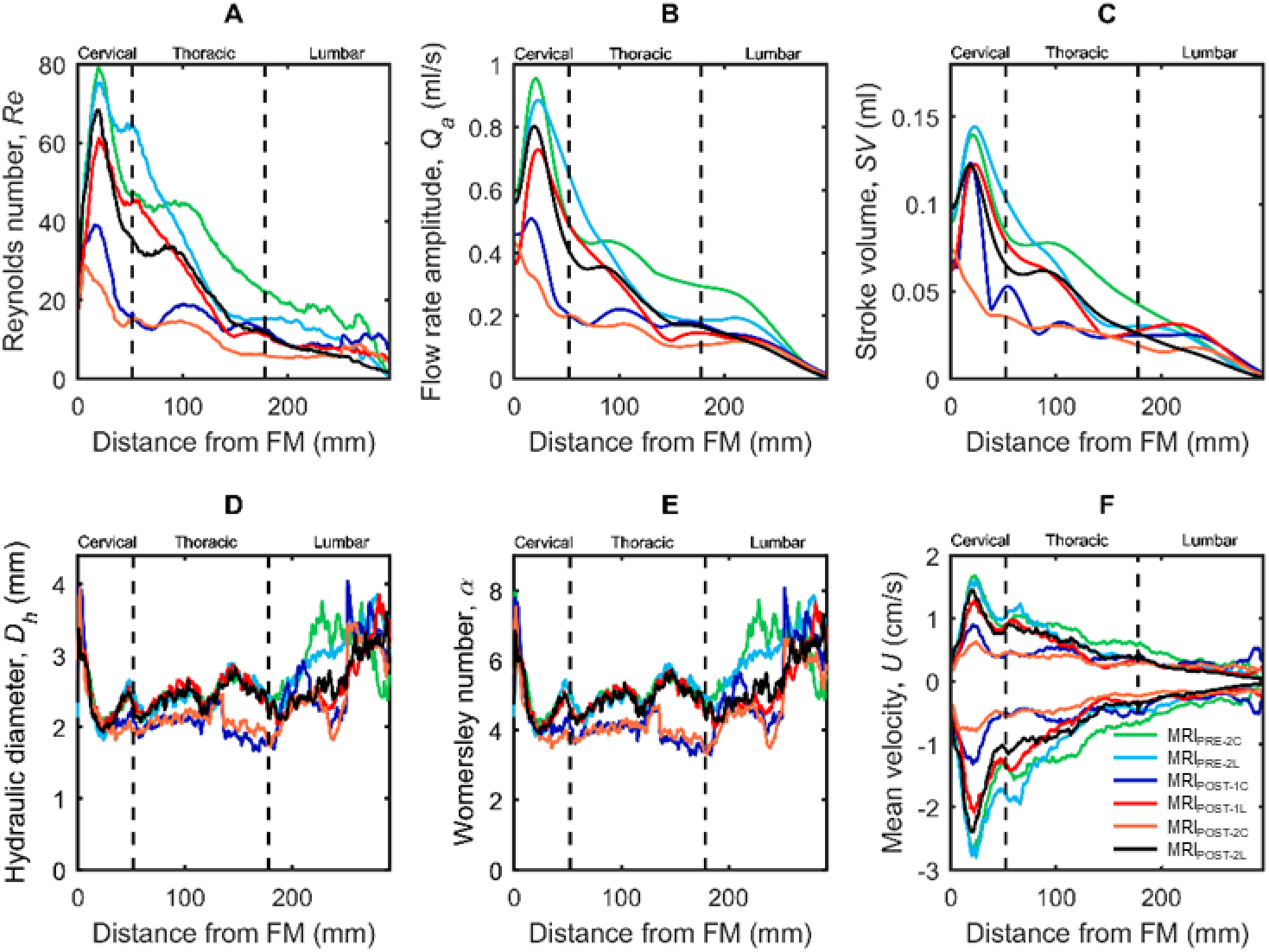
Hydrodynamic parameter axial distribution computed along the spine for cynomolgus monkeys. (A) Reynolds number, *Re*, (B) Flow rate amplitude, *Q*_*a*_, (C) Stroke Volume, *SV*, (D) left axis, Hydraulic diameter, *D*_*h*_, right axis, Womersley number, *α*, (E) mean peak systolic, 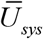, and diastolic, 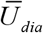, CSF velocity. Each line corresponds to mean value of each NHPs group with catheter located in the lumbar or cervical spine before or after catheter placement.

*D*_*h*_ (**Fig 6D**) and α (**Fig 6E**) decreased a great degree with cervical catheter implantation and to a lesser degree with lumbar implantation. Maximum *D*_*h*_ and α was 4 and 8 located near the FM. The peak value of the mean velocity ranged from +1.8 to −2.9 cm/s in MRI_PRE-2_ and occurred at the C3-C4 level (**Fig 6F**). Based on Hagen–Poiseuille equation, CSF flow reduction was predicted to be 48% after cervical implantation and 6% after lumbar implantation. These predictions were comparable to the MRI-measured *Q*_*peak-sys*_ reduction of 55% after cervical implantation and 21% after lumbar implantation (**Table 2**).

## Discussion

To the best of our knowledge, the impact of intrathecal catheter implantation on spinal CSF dynamics in a cynomolgus monkey has not been reported in the literature. Our results show that catheter implantation decreases spinal CSF dynamics and that the decrease is greater for cervical implantation compared to lumbar implantation. Also, that the decrease in spinal CSF dynamics was present immediately post-implantation and persisted two weeks after implantation.

### Catheter insertion decreased spinal CSF flow

The potential impact of catheter implantation on intrathecal CSF dynamics should be considered when implanting spinal catheters in NHPs and potentially humans. Although catheter diameter is relatively small, our results showed that cervical catheter implantation reduced peak CSF flow by 54% compared to 21% for lumbar implantation (**Fig 4** and **5**). Additionally, nearly all measures of CSF dynamics were altered to a greater degree for cervical implantation compared to lumbar implantation (**Table 2**). These results were further supported by estimation of CSF flow reduction, based on the Hagen-Poiseuille equation, indicating that the CSF flow reduction was likely due to increased hydraulic resistance stemming from the catheter’s reduction in subarachnoid space hydraulic diameter (**Fig 3** and **Table 2**).

The reduction in CSF flow could also potentially be attributed to inflammatory and/or infection post-catheter insertion, as documented in previous research (24). However, given that a) the reduction in CSF flow remained weeks following catheter insertion, b) the magnitude of flow reduction agreed with the estimated reduction based on fluid physics, and c) CSF flow reduction was greater for cervical catheter insertion, we believe the most probable source of CSF flow reduction to be increased hydraulic resistance directly due to the catheter.

In combination, the results indicate that to preserve normative intrathecal CSF flow, catheter placement should be located within the shortest length of the spine as possible and / or catheter diameter should be minimized to reduce its potential impact on hydraulic resistance within the spinal subarachnoid space. However, a smaller diameter catheter may not allow infusion of a desired flow rate or could potentially produce presence of a flow jet near the catheter tip. These factors could be assessed by parametric simulations. Alternatively, if possible, lumbar puncture should be applied as it would have minimal impact on CSF hydraulic resistance within the spine in NHPs or humans. However, for prolonged intrathecal drug delivery applications, catheter insertion may be the only viable for drug delivery.

Average *PWV* was found to be 1.15 m/s across all NHPs and was not impacted by catheter implantation (**Fig 5**). This is a potential indicator that spinal compliance, and likely intracranial pressure, was not affected due to catheter implantation. CSF *PWV* was previously measured by our research group in NHPs and found to have a similar value at 1.13 m/s (19). However, in humans, CSF *PWV* was measured to be 1.94 m/s (25), indicating that *PWV* within the spine in humans to potentially be different than NHPs.

### Spinal NHP CSF dynamics were laminar and inertial dominated

CSF flow remained laminar throughout the CSF flow cycle for all cases analyzed. Results showed that CSF dynamics were affected the most in the cervical spine near the C5 vertebral level in MRI_POST-2C_ with a maximum *Re* of 28, 100% less than MRI_PRE-2C_ (**Fig 6**). *Re* was computed to represent the ratio of steady inertial forces to viscous forces and help indicate whether laminar flow (<2300) was present at each phase-contrast slice location (**Fig 2**). A laminar CSF flow indicates that the flow is smooth with relatively little lateral mixing. This is different from a turbulent flow, where chaotic changes in pressure and velocity occur and can lead to a large increase in lateral mixing. Chaotic CSF velocity or pressure fluctuations are not expected to occur before or after catheter placement. However, it is possible that disease states that result in strongly elevated CSF flow velocities (jets) could result in turbulence (26).

Inertial effects are expected to dominate the SAS CSF flow field for normal physiological flow rates, frequencies and CSF fluid properties. *α* varied along the spine in a similar fashion as *D*_*h*_ with a minimum and maximum value of 3.8 and 8.1 (**Fig 6**). *α* was computed to quantify the ratio of unsteady inertial forces to viscous forces that impact the CSF velocity profile shape (27). For *α*<2, the CSF velocity profiles will be parabolic in shape and considered quasi-static. For 2<*α*<10 velocity profiles will be M-shaped and, for *α*>10, velocity profiles will be relatively flat (plug shaped) (28). The maximum value of *α* in the thoracic region decreased to ~4 after cervical catheter insertion. This means that the CSF velocity profiles will have a M-shape throughout the spine. However, the upper cervical and lumbar spine had higher *α* indicating a relatively flat velocity profile within those regions. Our previous computational fluid dynamics NHP model without catheter implantation indicated a relatively blunt CSF velocity profile in the cervical spine (29). It is not possible to confirm if the in vivo velocity profiles measured in the current study were blunt shaped (**Fig _2C_**) as the MRI resolution was not fine enough to accurately capture the relatively thin boundary layer expected in a blunt or M-shaped flow profile.

### Potential relevance of results with respect to intrathecal drug delivery

Based on the statistical analysis, catheter implantation led to decreased CSF flow rate within the spinal SAS, most notably under cervical implantation. In principle, a lower CSF flow rate is expected to decrease solute transport in the spine. Thus, it is expected that cervical catheter implantation would decrease solute transport to a greater degree than lumbar implantation. However, previous research (our paper, ref), and our current study results (Re in **Fig 6A** and **Table 2**), indicate that CSF velocities and streaming in the cervical spine are much greater than the lumbar spine. Thus, although catheter placement in the cervical spine may result in decreased CSF flow, drug delivery in this region may still allow more rapid mixing compared to the lumbar spine. Catheter implantation location may also need to be taken into account alongside potentially diminished CSF flow dynamics in disease states, such as ALS (30). Optimal catheter implantation location can be explored in future work in combination with the potential role of catheter implantation on CSF flow dynamics, but was outside the scope of the present research.

Based on the results, it can be hypothesized that the impact of catheter implantation on CSF dynamics would potentially be greater in cynomolgus monkeys compared to adult humans due to relatively smaller SAS cross-sectional area in NHPs compared to humans (10X greater (19, 30)). The average catheter diameter of 1.5 mm used in this study for cynomolgus monkeys is within the range of catheter diameters used in humans, ranging from 1.2 to 1.65 mm in outer diameter (31). Given the relatively smaller catheter diameter applied in adult humans, the potential impact of catheter implantation on CSF flow dynamics in adult humans may be relatively small. However, greater potential for catheter impact on CSF flow dynamics may be present in pediatric humans due to their relatively smaller SAS cross-sectional area compared to adults.

### Limitations and future directions

This study provides quantitative measures and comparison to investigate the impact of catheter insertion on intrathecal CSF dynamics and geometry in cynomolgus monkeys. Further studies should quantify the potential variance of these parameters in a larger study size across NHP species, age, sex, weight, and in disease states. Geometric characterization did not take into account spinal cord nerve root surface area or volume, which may account for ~231 cm^2^ and ~6 ml, respectively within the SAS in humans (32). It is expected that these structures will alter the SAS surface area results presented in the current study. Albeit, the surface area in contact with the spinal cord and dura is likely similar since the junction of spinal cord nerve roots with these structures is relatively small. Also, we do not expect these structures to alter spinal cord and dura surface area to a great degree or total SAS volume.

There are also a few unknowns in relation CSF flow dynamics. First, CSF flow coupling with the cardiovascular cycle is accounted for in the present study. However, CSF flow is also affected by respiration (33), which was not considered in this study using cardiac-gated PC-MRI measurements. Future studies could investigate the relative contribution of respiration and cardiovascular pulsations to CSF flow dynamics along the spinal axis. Finally, CSF flow was measured at six axial locations and interpolated to generate a smooth distribution along the spine. The ideal study would minimize or eliminate interpolation as much as possible by adding more axial slice locations. Also, CSF dynamics should be quantified within the intracranial space to better understand the exact distribution of CSF flow disruption that a spinal catheter may produce. However, in the present study, MRI time limitation for each NHP did not allow additional slice measurement locations. The focus of the present study was on the intrathecal space, as this region is most nearby intrathecal therapeutic injection location that can be accessed by lumbar puncture or other relatively minimally invasive procedures. Injection of medications within the ventricular space of the brain or cortical SAS would also be impacted by nearby CSF dynamics within the ventricles and cisterns of the brain.

The axial distribution for all geometric parameters tended to have a similar trend (**Fig 3**) indicating a strong dependence among geometric parameters. This means that if one parameter shows a significant difference between two conditions or two-time points, some of the other parameters should also display a significant difference. On the other hand, if only one parameter shows a significant difference, such significance may be due to experimental error and may not be reliable. Therefore, although nine of the 42 p values in Table 3 are significant, they are not consistent with the dependence among the parameters and therefore should be interpreted with caution.

## Conclusions

This study presents a detailed geometric and hydrodynamic characterization of intrathecal CSF dynamics for eight cynomolgus monkey (Macaca fascicularis) to quantify the differences that occur based on catheter placement location in the cervical compared to the lumbar spine. The overall findings were: 1) Catheter insertion decreases CSF dynamics within the spine, 2) These changes in CSF dynamics were greater for cervical implantation compared to lumbar catheter implantation, and 3) The decreases in CSF dynamics persisted up to two weeks post-catheter implantation. In combination, these results support that intrathecal catheter implantation can adversely impact CSF flow dynamics in the spinal SAS.

## Supplementary files

**S1 Table. Source data for the axial distribution of SAS geometric and hydrodynamic parameters and the CSF flow waveforms collected at different vertebral levels.** Data for all eight NHPs measured before catheter implantation (MRI_PRE-2_) and after catheter implantation (MRIPOST-1 and MRIPOST-2).

## Acknowledgements

None.

## Data availability

All relevant data are within the manuscript and its Supporting Information files.

